# Asian elephants (*Elephas maximus*) recognise human visual attention from body and face orientation

**DOI:** 10.1101/2025.04.27.650167

**Authors:** Hoi-Lam Jim, Shinya Yamamoto, Pakkanut Bansiddhi, Joshua M. Plotnik

## Abstract

Visual attention has mostly been studied in primarily visual species, such as dogs and nonhuman primates. Although elephants rely more on acoustic and olfactory cues, they also use visual displays and gestures to communicate. Smet and Byrne (2014) showed that African savanna elephants (*Loxodonta africana*) recognise human visual attention based on face and body orientation, but this has not been investigated in Asian elephants (*Elephas maximus*). Here, we examined whether Asian elephants are sensitive to human attentional states. We tested ten captive female elephants in Thailand and analysed the frequency of experimenter-directed signals in a food-requesting task based on the experimenter’s body and face orientation. Elephants gestured significantly more when the experimenter’s body was oriented towards them than when it was turned away, and they gestured most frequently when her face was also oriented towards them. This suggests that body orientation may be the more salient visual cue, with face orientation contributing to the effect only when accompanied by body orientation. These findings show that Asian elephants, like African savanna elephants, understand the importance of visual attention for effective communication, contributing to our understanding of cognitive abilities across the elephant taxon and visual attention in animals.

## 1. Introduction

Communication can be defined as the transfer of information from a signaller to a recipient, where the signal conveys information that may influence the behaviour of the receiver (regardless of whether they were the intended recipient) or both participants [1]. Importantly, the signal must be detected and perceived by the recipient for effective communication [2]. Animals can use a variety of signal modalities for communication, such as visual, auditory, or olfactory, and signals can be multimodal. Different species have adapted to use different senses to perceive and interpret the world around them to help them survive in their environment. For example, animals that have evolved a sophisticated visual system, including humans and nonhuman primates [3,4], often use visual signals to communicate with each other and use vision to navigate their social world [5]. Studies have shown that great apes adjust their communicative signals depending on the recipient’s attentional state. Chimpanzees (*Pan troglodytes*), for example, used more visual gestures when an experimenter faced them and more vocalisations when she faced away [6]. Chimpanzees, bonobos (*Pan paniscus*), and orangutans (*Pongo pygmaeus*) produced more begging behaviours (i.e., visual and/or auditory signals) when both the experimenter’s face and body were oriented towards them [7]. Similarly, orangutans and gorillas (*Gorilla gorilla gorilla*) used more visual gestures when the experimenter was facing them but did not vocalise more when she faced away [8].

Previous studies have primarily focused on apes and other visual animals, such as dogs. However, the evolutionary origins of this communicative ability are not well understood, particularly in non-primate, non-visual dominant species. To better understand the factors that have shaped sensitivity to visual communicative signals, it is important to expand research to a broader range of species. Elephants, with their highly social nature and reliance on audition and olfaction over vision— evidenced by the significantly larger brain areas dedicated to these senses compared to the visual cortex [4,9]—are an ideal species for this purpose. However, they still use visual displays and gestures to communicate with one another [10], which may be particularly beneficial in their complex fission-fusion societies, where cooperation is essential [11,12]. For example, Smet and Byrne [13] suggested that African savanna elephants (*Loxodonta africana*) use the “periscope-sniff” as an ostensive pointing signal, potentially helping others detect and respond to dangers.

Additionally, a study on greeting behaviour in African savanna elephants found that they adjust their gestures based on the recipient’s visual attention, preferring visual gestures when the recipient was looking and tactile gestures when they were not [14]. Their sensitivity to the attentional states of others may extend to humans as well: Smet and Byrne [15] found that captive elephants modified their experimenter-directed signals in a food-requesting task based on the experimenter’s face and body orientation, gesturing more when her face was oriented toward them—but only when her body was also turned sideways or toward them—consistent with Kaminski *et al*.’s [7] findings in apes. It is important to note that the animals in these studies had extensive experience with humans, which could make them more sensitive to human attentional states.

Despite these recent advances in our understanding of African savanna elephants’ socio-cognitive abilities, little is known about this in other elephant species. The three living elephant species—African savanna, African forest (*Loxodonta cyclotis*) and Asian (*Elephas maximus*)—belong to the family Elephantidae, with the lineage of Asian elephants diverging from that of African elephants approximately 5-7 million years ago [16]. Over this timespan, significant species-level differences in behaviour and ecology may have emerged [8,17,18], raising questions about the extent to which cognitive capacities are shared across the elephant taxon. Furthermore, although evolutionary rates may differ, the split between *Elephas* and *Loxodonta* is comparable in timing to the divergence between *Homo* and *Pan* lineages in the family Hominidae [16,19]. Several global environmental changes were occurring at that time, potentially affecting many lineages and contributing to speciation across diverse mammalian groups [16]. Studying these closely related species may therefore provide insights into how socio-cognitive abilities evolved under different ecological and social pressures. For example, Asian elephants [20,21] and African savanna elephants [22] performed similarly on a means-end task, suggesting comparable problem-solving abilities. However, other research has found differences—African savanna elephants can use human pointing cues to locate hidden food [23,24], but this ability has not been demonstrated in Asian elephants [9,25].

To further investigate potential cognitive similarities between elephant species, we tested Asian elephants’ understanding of human visual attention, closely following the methodology of Smet and Byrne [15]. Thus, we examined whether Asian elephants adjust the frequency of experimenter-directed signals in a food-requesting task based on the experimenter’s body and/or face orientation. For effective visual communication, the signaller must ensure they are within the recipient’s line of sight. Therefore, we predicted that elephants would gesture more frequently when the experimenter was present compared to when she was absent, and when the experimenter’s face and body was oriented towards them compared to when either was oriented away. Since the surface area of the human body is much larger than that of the face, and elephants’ visual acuity may not be sufficient to detect whether a human’s face is oriented towards or away from them at a distance, we further predicted that body orientation would be a stronger visual cue than face orientation.

## 2. Methods

Our hypotheses, predictions, study design, and the behavioural and statistical analysis plan were pre-registered (https://aspredicted.org/J32_HR5). We made two deviations from this: the elephants were tested in four sessions instead of two, and we modified Smet and Byrne’s [15] ethogram.

### Participants

Ten female captive Asian elephants (*Elephas maximus*) aged 11–61 (*M* = 36.8, *SD* = 17.16) from the Golden Triangle Asian Elephant Foundation living on the properties of the Anantara Golden Triangle Elephant Camp and Resort in Chiang Rai, Thailand, participated in the experiment between February and March 2024 (electronic supplementary material, table S1).

### Experimental design

Each elephant was tested in four sessions conducted on separate days using a within-subjects, repeated-measures design. There was a 2-day break between sessions 1 and 2, a 6-day break between sessions 2 and 3, and a 2-day break between sessions 3 and 4.

In each session, the elephant completed one trial of each of the following conditions, which varied the orientation of the experimenter’s body and head, in a randomised order (figure 1):

**Figure 1.**
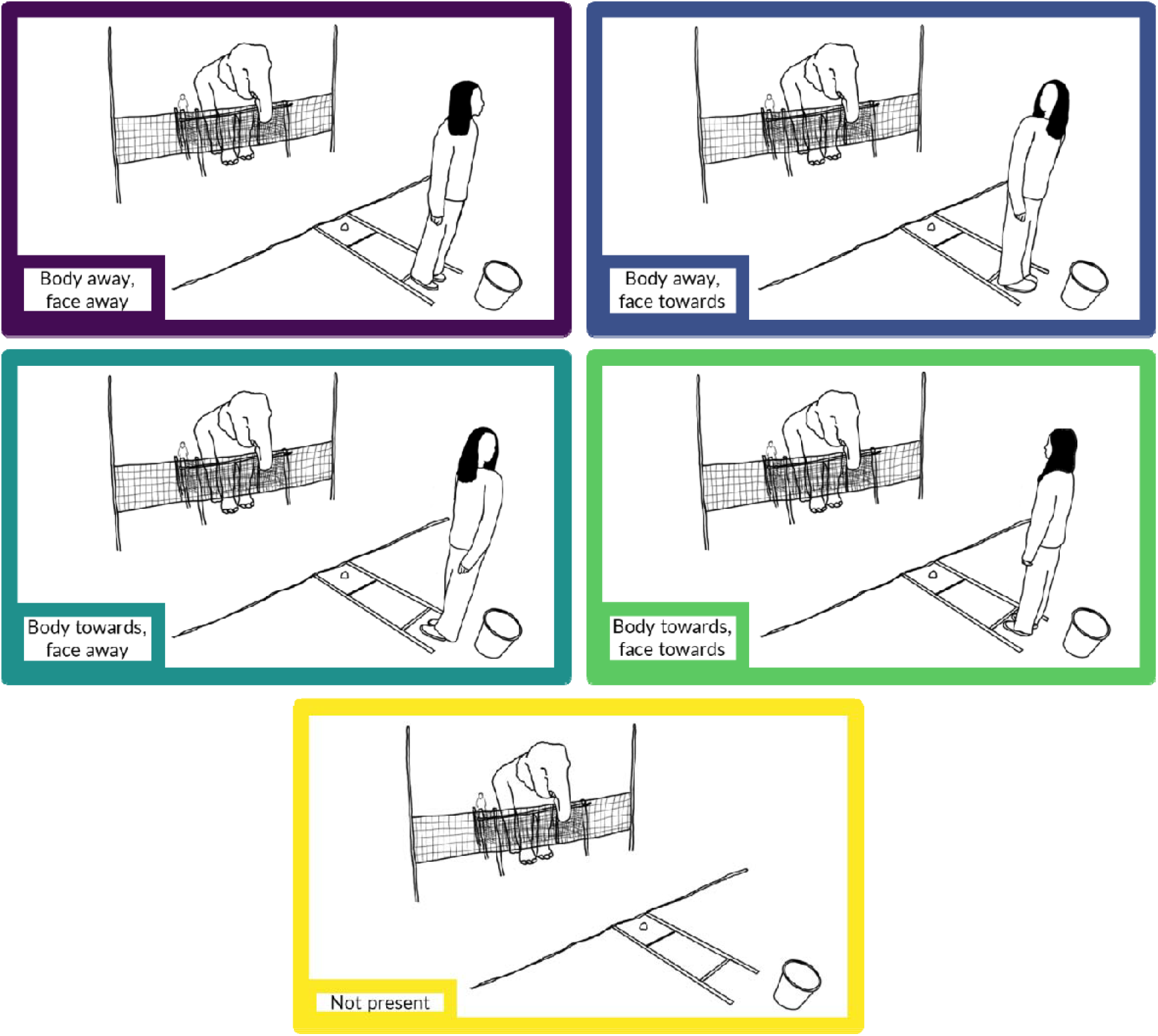
Experimental conditions showing the different orientations of the experimenter’s body and head, and the baseline condition. Illustrations by Hoi-Lam Jim.

1. Body away, face away (Ba_Fa)
2. Body away, face towards (Ba_Ft)
3. Body towards, face away (Bt_Fa)
4. Body towards, face towards (Bt_Ft)
5. Not present (Np) – henceforth ‘baseline’

We did not include the ‘body sideways’ conditions tested in Smet and Byrne [15] due to time constraints.

### Experimental setup

The experiment was conducted in a large field at the Anantara Golden Triangle Elephant Camp and Resort. The grass was cut in the testing area (10.6 × 21.5 m) and there was tall grass beyond this area. A 4.7 (L) × 1.2 m (H) volleyball net was strung in the centre of the field. A 2 (W) × 3.7 m (L) holding pen was built 35 cm behind the volleyball net to aid the elephants’ position during the experiment. The front and sides of the holding pen were covered with a volleyball net and the elephant was free to leave the holding pen from the back. The experimenter (H.-L.J., hereafter ‘E’) used a wooden tray, which was 50 × 50 cm with 1 m long handles, to feed the elephant during the experiment.

The whole experiment was recorded by two GoPro Hero 10 Black cameras. The side view camera was placed on a tripod at the side of the testing area and the front view camera was placed on a tripod close to E, facing the participant (figure 2).

**Figure 2.**
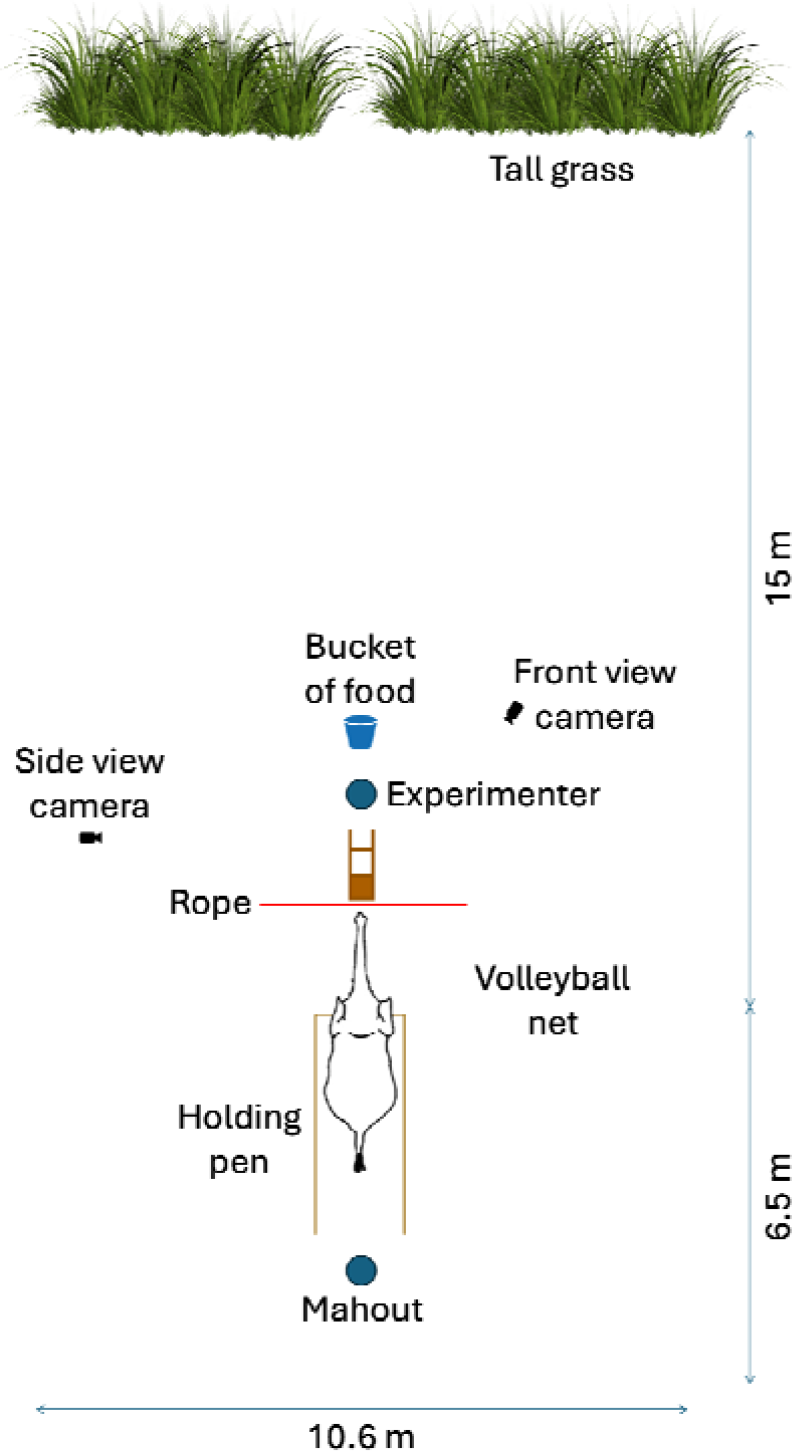
Schematic depiction of the experimental setup.

### Procedure

Wild grasses were available *ad libitum* outside experiment times, and the elephants were additionally fed according to their regular feeding regimes; thus, they were not food-deprived before the experiment. Mangoes, a high-value food reward for elephants, were cut into half-pieces and used as the food reward. Each elephant was accompanied by her own mahout (caretaker), who stood behind her during the experiment to avoid influencing her behaviour. However, there was one exception to this: due to the potential danger of working with one elephant, Yuki, her mahout wore sunglasses and initially stood to the side of her with his back towards E, so he was unaware of E’s posture. Once the mahout felt that Yuki was comfortable and the situation was safe, he moved to stand behind her. Throughout the experiment, the mahout only spoke to give commands, such as instructing the elephant to take the food if she did not reach for it when offered, or to stop if she physically interacted with the holding pen or volleyball net, or attempted to walk around the pen to retrieve the food.

Prior to testing, the elephant could explore the environment freely for approximately 5 minutes to familiarise herself with the location and the holding pen. Each mahout tested how far his elephant could reach with her trunk inside the holding pen, which ranged from 2.35–3.5 m, and the distance was marked by placing a rope on the ground. The tray was placed behind the rope; thus, the elephant supposedly could not reach the food. However, there were two occasions where the elephant managed to retrieve the food herself: one elephant, Jathong, pushed forward inside the holding pen, grabbed the tray, and pulled it towards herself; another elephant, Bo, retreated from the holding pen and walked around the volleyball net to eat from the tray and her mahout could not stop her in time. Both incidents occurred during the baseline after the test trial ended, thus the data were not excluded from analysis.

The experiment generally followed the procedure outlined in Smet and Byrne [15]. All elephants were tested individually between 7:30-9 am or 2-3 pm depending on their availability, and each session took approximately 10 minutes. A session began with three ‘no-delay’ trials: E stood behind a wooden tray, called the elephant’s name whilst facing her, and placed a piece of food onto the tray. E then immediately picked up the tray and moved forward to allow the elephant to eat from the tray. After the elephant took the food, E placed the tray down in its original position. If the elephant did not take the food from the tray voluntarily in the first ‘no-delay’ trial because she was afraid of touching the volleyball net, the mahout showed the elephant the food and encouraged her to take it from the tray, and then another ‘no-delay’ trial was conducted to ensure the elephant was comfortable with taking the food from the tray by herself. Elephants also had additional ‘no-delay’ trials during the experiment if a brief interruption or minor experimental mistake occurred. After three consecutive ‘no-delay’ trials, the testing phase began with the first test trial.

In the test trial, E stood behind the tray, called the elephant’s name, placed the food on the tray, picked it up to show it to the elephant, and put it down without giving the food. Then, E adopted one of the four postures (or walked away in the baseline), started the stopwatch, and stood still for 20 seconds (i.e., the test trial period) before picking the tray up again and moving forward to feed the participant. In the baseline, E started the stopwatch as she turned to walk quickly to the tall grass and hid behind it; thus, she was obscured from the elephant’s view and ‘not present’. After 20 seconds passed, E walked back to her starting position and moved the tray within the elephant’s reach (electronic supplementary material, video S1). Each test trial alternated with a ‘no-delay’ trial and sessions always ended with a ‘no-delay’ trial (see electronic supplementary material, figure S1 for a flowchart of the procedure).

### Behavioural analysis

We coded the elephants’ actions towards E and food during the test trial from the footage from the camera facing the elephant, and we supplemented it with the side view video footage when necessary.

We coded the frequency of head and trunk gestures produced to request food based on Smet and Byrne’s [15] ethogram. Some modifications were made because the elephants performed trained begging behaviours not observed in the previous study, and we refined the definitions by introducing subcodes (table 1). Each behaviour was coded as a single event (i.e., if a behaviour was performed three times consecutively, it was coded as three separate behaviours).

**Table 1.** Definitions of coded behaviours.

| Behaviour | Definition |
| --- | --- |
| Trunk swing | Tossing the trunk in the direction of E and/or the food in a quick, swift motion. |
| Trunk out | Slowly moving or holding some part of the trunk in the direction of E and/or the food. There were three subcodes based on the direction of the trunk tip: (1) forward (towards E); (2) self (towards the trunk or body); and (3) ground (towards the food). |
| Trunk up | Trunk upwards in an S-shape or holding some part of the trunk high above the head. There were three subcodes based on the direction of the trunk tip: (1) forward (towards E); (2) self (towards or touching the trunk); and (3) to the side. |
| Reach out | Full extension of the trunk. There were two subcodes based on where the trunk was pointing towards: (1) E or (2) the food. |
| Head nod | Head bobbing up and down at least once. One bout was coded as one event. |
| Trained begging | An “unnatural” behaviour taught by a human that is consistently performed on a verbal command. Only applicable to two elephants who performed distinct behaviours (Bo nodded her head and vocalised simultaneously, and Benz held her trunk out towards E, closed the airway to her trunk by holding the tip tightly together, then released it, creating a soft, suction-like sound). |

### Statistical analysis

Following the pre-registered analysis, we fitted a zero-inflated Poisson generalised linear mixed-effect model (GLMM) using the *glmmTMB* package (v1.1.9, [26]) to examine the frequency of head and trunk gestures in response to E’s visual attention. The test predictor was ‘condition’ (factor with five levels), with ‘session’ and ‘trial’ as z-transformed continuous covariates. Participant ID was included as a random effect. Pairwise comparisons with Tukey correction were conducted using the *emmeans* package (v1.10.2, [27]). For more information, see electronic supplementary material, methods.

## 3. Results

The GLMM showed that the frequency of head and trunk gestures elephants produced to request food varied significantly across experimental conditions (full-null model comparison: χ^2^ = 32.333, *df* = 4, *p* < .001; table S2). Gesture frequency significantly decreased across sessions (estimate = -0.318, *SE* = 0.078, χ^2^ = 16.571, *df* = 1, *p* < .001), possibly due to fatigue or learning effects—since E always provided food after trials, elephants may have learned that gesturing was unnecessary to receive the reward. Gesture frequency did not change significantly across trials (χ^2^ = 0.070, *df* = 1, *p* = .791).

To simplify the presentation of pairwise comparisons, the experimental conditions are abbreviated as in 2(c) (Experimental design). Elephants gestured significantly more in the two conditions where E’s body was oriented towards them compared to the baseline (Np – Bt_Fa: *p* = .003; Np – Bt_Ft: *p* < .001). In contrast, there was no significant difference in gesture frequency between the baseline and the two conditions where E’s body was oriented away (Np – Ba_Fa: *p* = .108; Np – Ba_Ft: *p* = .058). These results indicate that the mere presence of a human did not increase gesturing, and that elephants were sensitive to body orientation.

When body orientation was held constant, face orientation did not significantly affect gesture frequency, whether E’s body was oriented away (Ba_Fa – Ba_Ft: *p* = .998) or towards the elephants (Bt_Fa – Bt_Ft: *p* = .335). When face orientation was held constant, E’s body orientation had no significant effect when her face was oriented away (Ba_Fa – Bt_Fa: *p* = .674). However, elephants gestured significantly more when E’s body was oriented towards them, but only when her face was also oriented towards them (Ba_Ft – Bt_Ft: *p* = .042). These findings suggest that body orientation is the stronger cue overall, and its effect was enhanced when E’s face was also oriented towards the elephants. This is further supported by the stronger effect observed when both E’s body and face were oriented towards the elephants compared to when both were oriented away (Ba_Fa – Bt_Ft: *p* = .013), which was greater than the effect of Ba_Ft – Bt_Ft (*p* = .042) (figure 3, electronic supplementary material, table S3).

**Figure 3.**
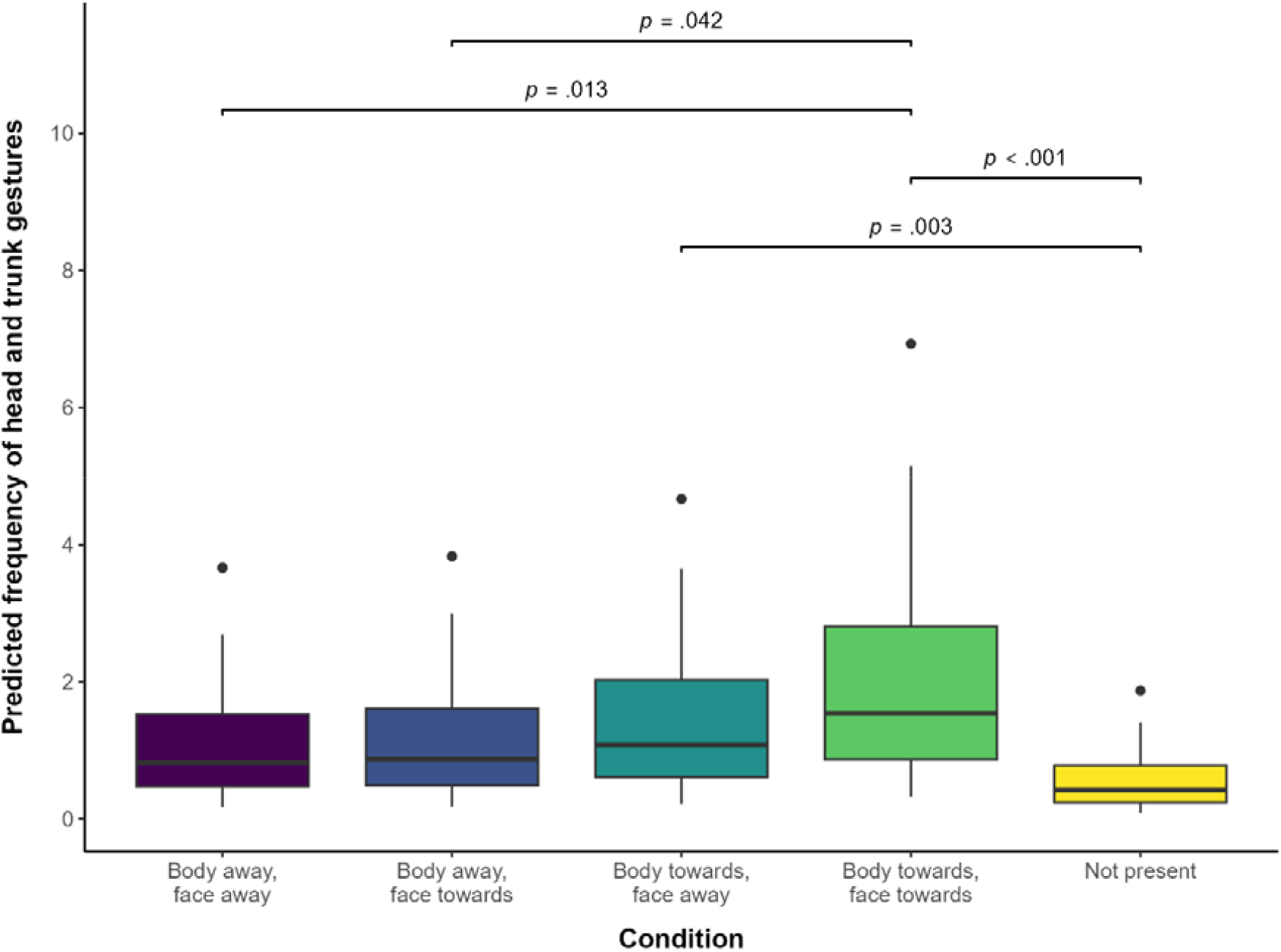
Box plot showing the predicted frequency of elephants’ head and trunk gestures when requesting food across experimental conditions. Black dots indicate outliers.

## 4. Discussion

Our results support the hypothesis that Asian elephants are sensitive to the attentional state of others and use visual gestures as communicative signals. Specifically, they performed more experimenter-directed signals in a food-requesting task when the experimenter’s face and body was oriented towards them compared to when either was oriented away. However, they did not gesture more merely because a human was present compared to when she was absent. Although Kaminski *et al*. [7] and Smet and Byrne [15] concluded that apes and African savanna elephants adjust their visual signals based on audience presence, they also found no significant difference in gesture frequency between the ‘body away, face away’ and ‘experimenter absent’ conditions. Thus, our findings align with their own, suggesting that the presence of a human alone is not sufficient to elicit gesturing, but rather apes and elephants are specifically attuned to visual attentional cues.

Elephants gestured significantly more when the experimenter’s body was oriented towards them compared to away, and they gestured most when her face was also oriented towards them. These results are in line with previous studies on apes [6-8] and African savanna elephants [15] and supports Kaminski *et al*.’s [7] hierarchical, bivariate interpretation, in which body and face orientation convey different types of information—specifically, body orientation signals the human’s disposition to give food, while face orientation indicates their ability to perceive a communicative signal.

Although faces are important for communication in primates, this may not apply to elephants, and there are several possible explanations why elephants may prioritise body orientation over face orientation as a stronger cue. First, captive elephants rely on humans for food, and since the experimenter could only provide a reward when her body was oriented towards them, elephants may have learned to associate body orientation with food availability. Second, although humans can feed elephants without directly facing them, feeding interactions typically involve direct attention and a frontal orientation for safety reasons. Third, body orientation may be a more salient visual cue than face orientation because the body’s larger surface area makes it easier to detect from a distance. This is further supported by the fact that Asian elephants [9,24] and African savanna elephants [22] appeared unable to use gaze cues alone to find food, suggesting that the human face may be insufficiently salient for elephants to perceive without another cue like the body.

Human body orientation may be a particularly relevant cue for wild elephants, as they likely do not approach humans closely enough to perceive their face orientation. They have been shown to discriminate between humans based on olfactory, visual, and auditory cues; African savanna elephants in Amboseli National Park, Kenya, reacted more fearfully to the scent of garments previously worn by a Maasai warrior compared to a similar aged man from the agricultural Kamba tribe, and they reacted more aggressively to red clothing typically worn by Maasai than to neutral white clothing [28]. Elephants also exhibited stronger defensive and investigative behaviours following playbacks of Maasai men’s voices than those of Maasai women or boys [29]. Although these responses were based on auditory cues, the associated visual characteristics of these groups (i.e., differences in body shape between men, women, and boys) may be perceivable to elephants from a distance. Thus, recognising human body orientation from afar could provide an evolutionary advantage, allowing elephants to assess potential threats more effectively.

This study adds to the literature suggesting that Asian elephants [20,21] and African savanna elephants [22] share similar cognitive abilities. However, previous studies focused on means-to-end problem-solving behaviour, which is not directly related to the present study. Our study is more closely aligned with research on following human pointing cues, as both involve visual cue perception, but conflicting results have been found between the two species [9,23-25]. We kept our methodology as consistent as possible with Smet and Byrne’s [15], and the elephants in both studies had similar rearing histories, being born and raised in captivity and accustomed to close human interaction. Thus, further research is needed to determine whether discrepancies in elephants’ sociocognitive abilities reflect ecological differences or are influenced by captivity and human exposure [9].

The STRANGE framework was established to identify sampling biases in animal behaviour studies that may affect the reproducibility and generalisability of findings [30,31], including rearing history as described above. Some elephants in this study had prior experience with cognitive research, while others participated in another experiment for the first time simultaneously (electronic supplementary material, table S1). Additionally, we only tested elephants that were available and with whom it was safe to work. Like previous studies on apes [6-8] and elephants [15] investigating sensitivity to human attentional states, our study involved enculturated individuals with extensive human experience, which may limit the ecological validity of the results. Finally, our small sample size means that individual variation in behaviour and cognition could affect the generalisability of our results.

In conclusion, this study is the first to demonstrate that Asian elephants are sensitive to human attentional states, prioritising body orientation as a key visual cue. Face orientation only played a role when combined with body orientation, similar to African savanna elephants. This research contributes to the literature on elephant cognition and supports the growing evidence for congruence between behavioural patterns in elephants and great apes, suggesting an underlying similarity in socio-cognitive mechanisms driven by convergent evolution.

## Supporting information

Data repository

## Ethics

This study was approved by the National Research Council of Thailand (Protocol #0401/95). Ethical approval was obtained from the Faculty of Veterinary Medicine’s Animal Care and Use Committee (Protocol #R23/2566) at Chiang Mai University and the Wildlife Research Center’s Ethical Committee (Protocol #WRC-2023-009A) at Kyoto University. The elephants’ participation was voluntary, and the mahout (caretaker) could stop the experiment at any time if he felt the elephant did not want to participate anymore, but this never happened.

## Data accessibility

The behavioural data and analysis code are available as electronic supplementary material and will be archived in an open data repository upon formal publication.

## AI declaration

We have not used AI-assisted technologies in creating this article.

## Authors’ contributions

H.-L.J.: Conceptualization, Data curation, Formal analysis, Funding acquisition, Investigation, Methodology, Project administration, Validation, Visualization, Writing – original draft, Writing – review & editing; S.Y.: Conceptualization, Funding acquisition, Methodology, Resources, Supervision, Writing – review & editing; P.B.: Project administration, Writing – review & editing; J.M.P.: Conceptualization, Methodology, Resources, Supervision, Writing – review & editing.

All authors gave final approval for publication and agreed to be held accountable for the work performed therein.

## Acknowledgements

We thank John Roberts, Poonyawee Srisantear, Nissa Mututanont, Nuttamon Chaichana, and the mahouts and staff of the Golden Triangle Asian Elephant Foundation for their dedication to the care and well-being of the elephants. We also thank Thitiphong Chintanawong and Nachiketha Sharma for their assistance with the study. We are grateful to the staff of Anantara Golden Triangle Elephant Camp and Resort for their support throughout our research. We also thank the National Research Council of Thailand for granting permission to conduct this study. Finally, we thank Sofia Vilela for interobserver reliability coding and Mayte Martínez for statistical guidance.

