## Supplementary Materials for "Asian elephants (*Elephas maximus*) recognise human visual attention from body and face orientation"

H.-L.J.: 0000-0002-8590-7167

S.Y.: 0000-0002-7556-6151

P.B.: 0000-0001-6699-4819

J.M.P.: 0000-0002-7597-8818

**Supplementary methods**

All statistical analyses were conducted using R (v4.4.1, R Core Team 2024) in RStudio (v2024.04.2+764, Posit team 2024). H.-L.J. and a research assistant independently coded 20% of the videos (10% from sessions 1 and 2 and 10% from sessions 3 and 4), which were randomly chosen. Both coders were blind to the condition in each test trial. Interobserver reliability was analysed using the Intraclass Correlation Coefficient (ICC) from the *irr* package (v0.84.1, Gamer et al. 2019) and revealed that inter-rater agreement was good (Koo and Li 2016) (ICC (two-way, agreement) = 0.88, $F = 15.4$, $p < .001$). H.-L.J. coded the remaining videos blind to the condition in each trial.

The model sample comprised 200 observations from 10 individuals, with 92 observations with no recorded behaviours. Since 46% of our data consisted of zeros, we applied a zero-inflated Poisson GLMM. Due to the limited data for individual behaviours, we combined all behaviours and used their total frequency as the response variable. To avoid multiple testing (Forstmeier and Schielzeth 2011) and keep type I error rate at the nominal level of 0.05, we compared the full model with a null model lacking the test predictor ('condition'). This comparison was tested by means of a likelihood ratio test using the *anova* function with the *method* argument set to "*Chisq*" (Dobson and Barnett 2018). The plot was generated using *ggplot2* (v3.5.1, Wickham 2016).

We checked model diagnostics as follows: collinearity was assessed by computing Variance Inflation Factors (VIF) (Field 2005) for a standard linear model using the *vif* function in the *car* package (v3.1-2, Fox and Weisberg 2019), which indicated no issues (maximum VIF = 1.014). Model stability was assessed by comparing the estimates obtained from the model based on all data with those obtained from models with the levels of random effects excluded one at a time, which revealed the model to be stable (see table S2 for estimate ranges). Confidence intervals were calculated using the *boot.glmmTMB* function. The model was not overdispersed (dispersion parameter = 1.147). R functions used for assessing model stability, confidence intervals and overdispersion were provided by Mundry (2023).

**Table S1.** Individual characteristics of participants.

| Elephant | Age (years) | Experience in other experiments |
|---|---|---|
| Bo | 46 | Yes – prior |
| Benz | 18 | Yes – first time simultaneously |
| Boonma | 61 | No |
| Boonrod | 29 | Yes – first time simultaneously |
| Boonsri | 56 | Yes – prior |
| Dah | 22 | Yes – prior |
| Jathong | 33 | Yes – prior |
| Kummool | 54 | Yes – prior |
| Yokfah | 11 | Yes – first time simultaneously |
| Yuki | 38 | Yes – prior |

**Table S2.** Results from the zero-inflated Poisson GLMM predicting the frequency of head and trunk gestures. Abbreviations: Ba_Fa = Body away, face away; Ba_Ft = Body away, face towards; Bt_Fa = Body towards, face away; Bt_Ft = Body towards, face towards; Np = Not present.

| Part[1] | Term | Estimate | SE | 95% CI | | Model stability | | *z* | *df* | *p*[2] |
|---|---|---|---|---|---|---|---|---|---|---|
| | | | | Lower | Upper | Min | Max | | | |
| Count | Intercept | -0.816 | 0.329 | -1.567 | -0.286 | -1.098 | -0.705 | | | |
| | Condition: Ba_Fa[3] | 0.658 | 0.271 | 0.175 | 1.247 | 0.539 | 0.739 | 2.427 | 4 | .015 |
| | Condition: Ba_Ft | 0.728 | 0.272 | 0.223 | 1.372 | 0.632 | 0.874 | 2.672 | | .008 |
| | Condition: Bt_Fa | 0.938 | 0.262 | 0.412 | 0.543 | 0.804 | 1.059 | 3.587 | | <.001 |
| | Condition: Bt_Ft | 1.282 | 0.249 | 0.828 | 1.901 | 1.128 | 1.444 | 5.152 | | <.001 |
| | Session[4] | -0.318 | 0.078 | -0.467 | -0.187 | -0.392 | -0.251 | -4.078 | 1 | <.001 |
| | Trial[4] | 0.018 | 0.068 | -0.129 | 0.163 | -0.017 | 0.055 | 0.265 | 1 | .791 |

| | | | | | | | |
|---|---|---|---|---|---|---|---|
| Zero | Intercept | -2.433 | 0.776 | -20.342 | -1.543 | -4.212 | -2.183 |

Estimate, standard errors (SE), 95% confidence intervals (CI), model stability (estimate ranges derived after excluding individuals one at a time) and results of significance tests (Wald's *z* approximation).

[1]'Count' indicates the count part and 'Zero' the zero-inflation part of the model

[2]The *p* value for the intercept is not shown due to its limited interpretability

[3]Reference level for Condition = Np

[4]Continuous variables (Session and Trial) were z-transformed (Session: *M* = 2.5, *SD* = 1.121; Trial: *M* = 3, *SD* = 1.418)

**Table S3.** Pairwise comparisons. Significant *p* values are in bold. Abbreviations: Ba_Fa = Body away, face away; Ba_Ft = Body away, face towards; Bt_Fa = Body towards, face away; Bt_Ft = Body towards, face towards; Np = Not present.

| Comparisons | Estimate | SE | *p* | 95% CI | |
|---|---|---|---|---|---|
| | | | | Lower | Upper |
| Np – Ba_Fa | -0.658 | 0.271 | .108 | -1.397 | 0.082 |
| Np – Ba_Ft | -0.728 | 0.272 | .058 | -1.471 | 0.015 |
| Np – Bt_Fa | -0.938 | 0.262 | **.003** | -1.651 | -0.225 |
| Np – Bt_Ft | -1.282 | 0.249 | **< .001** | -1.961 | -0.603 |
| Ba_Fa – Ba_Ft | -0.070 | 0.226 | .998 | -0.687 | 0.548 |
| Ba_Fa – Bt_Fa | -0.280 | 0.211 | .674 | -0.856 | 0.296 |
| Ba_Fa – Bt_Ft | -0.625 | 0.197 | **.013** | -1.162 | -0.087 |
| Ba_Ft – Bt_Fa | -0.210 | 0.214 | .864 | -0.795 | 0.374 |
| Ba_Ft – Bt_Ft | -0.555 | 0.199 | **.042** | -1.097 | -0.012 |
| Bt_Fa – Bt_Ft | -0.344 | 0.184 | .335 | -0.847 | 0.158 |

**Video S1.** Example of a test trial in the 'Body towards, face towards' condition.

https://youtu.be/6c3mXxT_SDA

**Figure S1.** Flowchart illustrating an example of the full procedure for one participant. Colours correspond to the experimental conditions shown in figures 2 and 3 of the main text. Abbreviations: Ba_Fa = Body away, face away; Ba_Ft = Body away, face towards; Bt_Fa = Body towards, face away; Bt_Ft = Body towards, face towards; Np = Not present.

| Session 1 | | Session 2 | | Session 3 | | Session 4 |
|---|---|---|---|---|---|---|
| 1. No-delay trial<br>2. No-delay trial<br>3. No-delay trial<br>4. Test trial 1 (Ba_Fa)<br>5. No-delay trial<br>6. Test trial 2 (Bt_Fa)<br>7. No-delay trial<br>8. Test trial 3 (Ba_Ft)<br>9. No-delay trial<br>10. Test trial 4 (Np)<br>11. No-delay trial<br>12. Test trial 5 (Bt_Ft)<br>13. No-delay trial | 2-day break | 1. No-delay trial<br>2. No-delay trial<br>3. No-delay trial<br>4. Test trial 1 (Np)<br>5. No-delay trial<br>6. Test trial 2 (Bt_Ft)<br>7. No-delay trial<br>8. Test trial 3 (Ba_Fa)<br>9. No-delay trial<br>10. Test trial 4 (Bt_Fa)<br>11. No-delay trial<br>12. Test trial 5 (Ba_Ft)<br>13. No-delay trial | 6-day break | 1. No-delay trial<br>2. No-delay trial<br>3. No-delay trial<br>4. Test trial 1 (Bt_Fa)<br>5. No-delay trial<br>6. Test trial 2 (Ba_Fa)<br>7. No-delay trial<br>8. Test trial 3 (Ba_Ft)<br>9. No-delay trial<br>10. Test trial 4 (Bt_Ft)<br>11. No-delay trial<br>12. Test trial 5 (Np)<br>13. No-delay trial | 2-day break | 1. No-delay trial<br>2. No-delay trial<br>3. No-delay trial<br>4. Test trial 1 (Ba_Ft)<br>5. No-delay trial<br>6. Test trial 2 (Np)<br>7. No-delay trial<br>8. Test trial 3 (Bt_Ft)<br>9. No-delay trial<br>10. Test trial 4 (Ba_Fa)<br>11. No-delay trial<br>12. Test trial 5 (Bt_Fa)<br>13. No-delay trial |
