## Supplementary material for "Asian elephants (*Elephas maximus*) recognise human visual attention from body and face orientation": Data repository: ele_vis_att_analysis.Rmd

````
R version 4.4.1 "Race for Your Life"

RStudio 2024.04.2+764 "Chocolate Cosmos"

```{r}
#R packages used
library(glmmTMB) #for zero-inflated negative binomial GLMM
library(emmeans) #for pairwise comparisons
library(magrittr) #for piping (%>%) operations
library(ggplot2) #for making graph
library(ggsignif) #for adding significance annotations
library(viridis) #for using colourblind-friendly colours in graph
library(car) #for VIF (collinearity)

#R functions for model assumptions by Mundry (2023)
source("../functions/diagnostic_fcns.r")
source("../functions/glmm_stability.r")
source("../functions/glmmTMB_stability.r")
source("../functions/boot_glmmTMB.r")
```

```{r}
xdata <- read.csv("../data/ele_vis_att_data.csv")

```

```{r}
hist(xdata$totalbehaviours, breaks = seq(min(xdata$totalbehaviours) - 0.5, 
                                         max(xdata$totalbehaviours) + 0.5, 
                                         by = 1),
     main = "Histogram of total behaviours",
     xlab = "Total behaviours", ylab = "Frequency",
     labels = TRUE)

#check how many 0s in data
sum(xdata$totalbehaviours == 0)
#check how many total behaviours
counts<-table(xdata$totalbehaviours)
counts_df<-as.data.frame(counts)
sum(counts_df$Freq)
92/200
```

because a large proportion of our data are zeros (46%), we will run a zero-inflated Poisson GLMM

z-transform trial and session because they are covariates

```{r}
mean(xdata$session)
sd(xdata$session)

mean(xdata$trial)
sd(xdata$trial)

xdata$z.session=as.vector(scale(xdata$session))
xdata$z.trial=as.vector(scale(xdata$trial))
```


```{r}
str(xdata)
xdata$condition <- as.factor(xdata$condition)
xdata$condition <- relevel(xdata$condition, ref = "Np")

full_zip=glmmTMB(totalbehaviours ~ condition + z.session + z.trial+(1|subject), ziformula = ~1, data = xdata, family=poisson)
full_zip$sdr$pdHess #model converged
overdisp.test(full_zip) #not overdispersed (1.147)

null_zip=glmmTMB(totalbehaviours ~ z.session + z.trial+(1|subject), ziformula = ~1, data = xdata, family=poisson)
null_zip$sdr$pdHess #model converged
overdisp.test(null_zip) #not overdispersed (1.365)

as.data.frame(anova(null_zip, full_zip, method="Chisq"))
as.data.frame(drop1(full_zip, test="Chisq"))
summary(full_zip)$coefficients

```

## Pairwise comparisons

```{r}
emm_zip <- emmeans(full_zip, ~ condition)
pairs(emm_zip)

pairs(emm_zip) %>% confint()
```

## Plot the model

```{r}
# Rename levels of the 'condition' variable
xdata$conditionsg <- factor(xdata$condition,
                          levels = c("Ba_Fa", "Ba_Ft", "Bt_Fa", "Bt_Ft", "Np"),
                          labels = c("Body away, face away", 
                                     "Body away, face towards", 
                                     "Body towards, face away", 
                                     "Body towards, face towards", 
                                     "Not present"))

#p value annotations
annotations = c("italic(p) == '.003'", 
                "italic(p) < '.001'", 
                "italic(p) == '.013'", 
                "italic(p) == '.042'")

# Define colors using viridis palette
colors <- viridis_pal()(5)

# Predict probabilities using the fitted model
xdata$predicted_prob_zip <- predict(full_zip, type = "response")

# Box plot by condition
ggplot(xdata, aes(x = conditionsg, y = predicted_prob_zip)) +
  geom_boxplot(fill = colors) +
  geom_signif(comparisons = list(
    c("Not present", "Body towards, face away"),
    c("Not present", "Body towards, face towards"),
    c("Body away, face away", "Body towards, face towards"),
    c("Body away, face towards", "Body towards, face towards")
  ),
  annotations = annotations,
    parse = TRUE,
  y_position = c(8, 9, 10, 11),
  tip_length = 0.01,
  textsize = 3.5
  ) +
  labs(x = "Condition", y = "Predicted frequency of head and trunk gestures") +
  scale_y_continuous(breaks = seq(0, 10, by = 2))+
  scale_x_discrete(labels = c('Body away,\nface away', 
                              'Body away,\nface towards', 
                              'Body towards,\nface away', 
                              'Body towards,\nface towards', 
                              'Not present')) +
  theme_classic() +
  theme(text = element_text(size = 12),  
        axis.title.x = element_text(margin = margin(t = 10), face = "bold"), 
        axis.title.y = element_text(margin = margin(r = 10), face = "bold"))
```

## Model assumptions

**collinearity (VIF)**

```{r}
xx_zip=lm(totalbehaviours~condition+z.session+z.trial, data = xdata) 
vif(xx_zip)
```

no collinearity

**Model stability**

```{r}
m.stab_zip=glmmTMB.stab(model.res=full_zip, para=T, data=xdata)
m.stab.plot(m.stab_zip$summary[, -1])
round(m.stab_zip$summary[,-1], 3) #min & max from the table
```

**BLUPs**

```{r}
ranef.diagn.plot(full_zip)
```

**confidence intervals (2.5% and 97.5%)**

```{r}
ci_zip=boot.glmmTMB(model.res=full_zip, data=xdata, excl.non.conv=F, nboots=1000, para=T, resol=100, level=0.95, use=NULL, contr=NULL, n.cores=c("all-1", "all"))
round(ci_zip$ci.estimates$fe, 3)
```
````
