## Supplementary material for "Asian elephants (*Elephas maximus*) recognise human visual attention from body and face orientation": Data repository: boot_glmmTMB.r

```
boot.glmmTMB<-function(model.res, data, excl.non.conv=F, nboots=1000, para=F, resol=100, level=0.95, use=NULL, contr=NULL, circ.var.name=NULL, circ.var=NULL, 
	n.cores=c("all-1", "all"), save.path=NULL, load.lib=T, lib.loc=.libPaths(), set.all.effects.2.zero=F){
	if(!summary(model.res)$link%in%c("log", "logit", "identity")){
		stop("link functions other than log, logit, or identity aren't supported yet; please contact Roger")
	}
	if(load.lib){library(lme4, lib.loc=lib.loc)}
	print("doesn't account for circular variables in the zero-inflated model")
	print("doesn't consider the dispersion part in the bootstrap of fitted values")
	print("haven't tested it yet with a cbind response")
	print("weights aren't supported yet")
	n.cores=n.cores[1]
	if(is.null(contr)){
		contr=glmmTMBControl()
	}
	##define function extracting all estimated coefficients (fixed and random effects) and also the model summary wrt random effects:
	extract.ranef.from.glmmTB<-function(m){
		#to.do=VarCorr(model.res)
		#varcor.names=names(to.do)
		to.do=summary(m)$varcor
		xx=lapply(to.do, function(x){
			lapply(x, attr, "stddev")
		})
		res=data.frame(
			what=rep(names(to.do), unlist(lapply(xx, function(x){sum(unlist(lapply(x, length)))}))),
			Groups=rep(unlist(lapply(to.do, names)), unlist(lapply(xx, function(x){unlist(lapply(x, length))}))),
			var1=unlist(lapply(xx, function(x){unlist(lapply(x, names))})),
			var2=NA,
			sdcor=unlist(xx)
		)
		xx=lapply(to.do, function(x){
			lapply(x, attr, "correlation")
		})
		xx=xx[unlist(lapply(xx, length))>0]
		xx=data.frame(
			what=rep(names(xx), unlist(sum(unlist(lapply(xx, function(x){lapply(x, function(x){prod(dim(x))})}))))),
			Groups=rep(unlist(lapply(to.do, names)), unlist(lapply(xx, function(x){lapply(x, function(x){prod(dim(x))})}))),
			var1=unlist(lapply(xx, function(x){lapply(x, function(x){rep(rownames(x), times=ncol(x))})})),
			var2=unlist(lapply(xx, function(x){lapply(x, function(x){rep(colnames(x), each=nrow(x))})})),
			sdcor=unlist(xx)
		)
		xx=xx[as.character(xx[, "var1"])<as.character(xx[, "var2"]), ]
		res=rbind(res, xx)
		res=res[order(as.character(res$what), as.character(res$Groups), as.character(res$var1)), ]
		return(res)
	}
	extract.BLUPs.from.glmmTB<-function(m){
		to.do=lapply(c("cond", "zi"), function(x){
			if(length(ranef(m)[[x]])>0){
			lapply(ranef(m)[[x]], function(y){
				return(list(
					what=rep(x, prod(dim(as.matrix(y)))),
					blup=unlist(c(as.matrix(y))),
					level=rep(rownames(y), times=ncol(y)),
					Name=rep(colnames(y), each=nrow(y))
				))
			})
			}else{
				return(list(list(what=NULL, blup=NULL, level=NULL, Name=NULL)))
			}
		})
		xc=sapply(lapply(to.do[[1]], "[[", "what"), length)
		xz=sapply(lapply(to.do[[2]], "[[", "what"), length)
		res=data.frame(
			what=as.vector(unlist(lapply(to.do, function(x){lapply(x, "[[", "what")}))),
			Grp=c(rep(names(xc), times=xc), rep(names(xz), times=xz)),
			Name=as.vector(unlist(lapply(to.do, function(x){lapply(x, "[[", "Name")}))),
			level=as.vector(unlist(lapply(to.do, function(x){lapply(x, "[[", "level")}))),
			blup=as.vector(unlist(lapply(to.do, function(x){lapply(x, "[[", "blup")})))
		)
		res=res[order(res$what, res$Grp, res$Name, res$level), ]
		return(res)
	}
	extract.all<-function(mres){
		##extract random effects model summary:
		vc.mat=extract.ranef.from.glmmTB(m=mres)
		##extract random effects, BLUPs:
		BLUPs=extract.BLUPs.from.glmmTB(m=mres)
		##dispersion parameter:
		dp=summary(mres)$sigma
		##fixed effects
		xx=fixef(mres)
		fe=data.frame(
			what=rep(names(xx), times=sapply(xx, length)),
			term=unlist(lapply(xx, names)),
			est=unlist(xx)
		)
		fe=fe[order(fe$what, fe$term), ]
		return(list(fe=fe, sigma=dp, vc.mat=vc.mat, BLUPs=BLUPs))
	}
	##prepare for new calls:
	xcall=as.character(model.res$call)
	names(xcall)=names(model.res$call)
	xcall=list(cond.form=xcall["formula"], zi.form=xcall["ziformula"], disp.form=xcall["dispformula"], xfam=xcall["family"])
	if(grepl(xcall[["xfam"]], pattern="(", fixed=T)){
		xfam=xcall[["xfam"]]
		xfam=unlist(strsplit(xfam, split="(", fixed=T))
		xfam[2]=gsub(x=xfam[2], pattern=")", replacement="", fixed=T)
		xfam[2]=gsub(x=xfam[2], pattern="link = ", replacement="", fixed=T)
		xfam[2]=gsub(x=xfam[2], pattern="\"", replacement="", fixed=T)
		if(substr(xcall[["xfam"]], start=1, stop=5)!="Gamma"){
			xcall[["xfam"]]=get(xfam[1])(xfam[2])
		}else{
			if(xfam[2]=="log"){
				xcall[["xfam"]]=Gamma(link="log")
			}else if(xfam[2]=="inverse"){
				xcall[["xfam"]]=Gamma(link="inverse")
			}else if(xfam[2]=="identity"){
				xcall[["xfam"]]=Gamma(link="identity")
			}else{
				stop("Error: family not supported")
			}
		}
	}
		
	xweights=try(model.res$frame[, "(weights)"], silent=T)
	if(class(xweights)[[1]]=="try-error"){
		xweights=rep(1, nrow(model.res$frame))
	}
	data$xweights.X.=xweights
	rv.name=gsub(unlist(strsplit(xcall[["cond.form"]], split="~"))[1], pattern=" ", replacement="", fixed=T)
	##define function doing the bootstrap:
	boot.fun<-function(x, xcall., data., rv.name., model.res., contr., excl.non.conv., save.path., extract.all.){
		xdone=F
		while(!xdone){
			done2=F
			while(!done2){
				data.[, rv.name.]=simulate(object=model.res.)[, 1]
				if(xcall[["xfam"]][["family"]]!="beta" | 
					(xcall[["xfam"]][["family"]]=="beta" & min(data.[, rv.name.])>0 & max(data.[, rv.name.])<1)){
					done2=T
				}
			}	
			i.res=glmmTMB(formula=as.formula(xcall.[["cond.form"]]), ziformula=as.formula(xcall.[["zi.form"]]), dispformula=as.formula(xcall.[["disp.form"]]), 
				data=data., control=contr., family=xcall[["xfam"]], weights=xweights.X.)
				if(i.res$sdr$pdHess | !excl.non.conv.){
					xdone=T
				}
		}
		conv=i.res$sdr$pdHess
		i.res=extract.all.(mres=i.res)
		if(length(save.path.)>0){save(file=paste(c(save.path., "/b_", x, ".RData"), collapse=""), list=c("i.res", "conv"))}
		return(list(boot.res=i.res, conv=conv))
	}
	##do bootstrap:
	if(para){
		library(parallel)
		cl <- makeCluster(getOption("cl.cores", detectCores()))
		if(n.cores!="all"){
			if(n.cores=="all-1"){n.cores=length(cl)-1}
			if(n.cores<length(cl)){
				cl=cl[1:n.cores]
			}
		}
		parLapply(cl=cl, 1:length(cl), fun=function(x, .lib.loc=lib.loc){
		  library(glmmTMB, lib.loc=.lib.loc)
		  return(invisible(""))
		})
		all.res=parLapply(cl=cl, X=1:nboots, fun=boot.fun, model.res.=model.res, xcall.=xcall, data.=data, rv.name.=rv.name, contr.=contr, 
			excl.non.conv.=excl.non.conv, save.path.=save.path, extract.all.=extract.all)
	}else{
    all.res=lapply(X=1:nboots, FUN=boot.fun, model.res.=model.res, xcall.=xcall, data.=data, rv.name.=rv.name, contr.=contr, 
			excl.non.conv.=excl.non.conv, save.path.=save.path, extract.all.=extract.all)
	}
	##extract results:
	##convergence:
	all.conv=sapply(all.res, "[[", "conv")
	##sigma:
	all.sigma=sapply(all.res, function(x){x$boot.res$sigma})
	##estimates, fixef effects:	
	all.fe=lapply(all.res, function(x){x$boot.res$fe$est})
	all.fe=matrix(unlist(all.fe), nrow=length(all.fe), byrow=T)
	colnames(all.fe)=apply(all.res[[1]]$boot.res$fe[, c("what", "term")], 1, paste, collapse="@")
	##estimates, random effects:
	all.re=lapply(all.res, function(x){x$boot.res$vc.mat$sdcor})
	all.re=matrix(unlist(all.re), nrow=length(all.re), byrow=T)
	colnames(all.re)=apply(all.res[[1]]$boot.res$vc.mat[, c("what", "Groups", "var1", "var2")], 1, paste, collapse="@")
	##estimates, BLUPs (well, it doesn't make any sense  to consider them):
	all.BLUPs=lapply(all.res, function(x){x$boot.res$BLUPs$blup})
	all.BLUPs=matrix(unlist(all.BLUPs), nrow=length(all.BLUPs), byrow=T)
	colnames(all.BLUPs)=apply(all.res[[1]]$boot.res$BLUPs[, c("what", "Grp", "Name", "level")], 1, paste, collapse="@")
	##store results:
	all.ind.boots=list(fe=all.fe, re=all.re, sigma=all.sigma, conv=all.conv)
	##get confidence intervals and merge them with original values:
	##fixed effects:
		orig=extract.all(model.res)
		xx=orig$fe
		xx$comb=apply(xx[, c("what", "term")], 1, paste, collapse="@")
		#xx$comb=paste(rownames(xx), xx$comb, sep="@")
		ci=apply(all.fe, 2, quantile, prob=c((1-level)/2, 1-(1-level)/2), na.rm=T)
		#times=unlist(lapply(summary(model.res)$coefficients, nrow))
		#comb=paste(rep(c("cond", "zi", "disp")[1:length(times)], times=times), unlist(lapply(summary(model.res)$coefficients, rownames)), sep="@")
		ci=ci[, match(xx$comb, colnames(ci))]
		#xx=xx[match(comb, xx$comb), ]
		ci.fe=data.frame(orig=xx$est, t(ci))
		##order according to original:
		ci.fe=ci.fe[paste(rep(c("cond", "zi", "disp"), times=unlist(lapply(fixef(model.res), length))), unlist(lapply(fixef(model.res), names)), sep="@"), ]
	##random effects:
		xx=orig$vc.mat
		xx$comb=apply(xx[, c("what", "Groups", "var1", "var1")], 1, paste, collapse="@")
		ci=apply(all.re, 2, quantile, prob=c((1-level)/2, 1-(1-level)/2), na.rm=T)
		ci.re=data.frame(orig=xx$sdcor, t(ci))
	##store results:
	ci.estimates=list(fe=ci.fe, re=ci.re)

	if(length(use)>0){
		ci.fitted.cond=NULL
		ci.fitted.zi=NULL
		ci.fitted.disp=NULL
		#extract fixed effects terms from the model:
		model.terms=c(attr(terms(as.formula(xcall[["cond.form"]])), "term.labels"), 
			attr(terms(as.formula(xcall[["zi.form"]])), "term.labels"), 
			attr(terms(as.formula(xcall[["disp.form"]])), "term.labels"))
		#exclude random effects:
		model.terms=model.terms[!grepl(x=model.terms, pattern="|", fixed=T)]
		#exclude random effects, interactions and squared terms from model.terms:
		model.terms=model.terms[!grepl(x=model.terms, pattern="I(", fixed=T)]
		model.terms=model.terms[!grepl(x=model.terms, pattern=":", fixed=T)]
		model.terms=unique(model.terms)
		
		#create new data to be used to determine fitted values:
		if(length(use)==0){use=model.terms}
		new.data=vector("list", length(model.terms))
		if(length(circ.var.name)==1){
			set.circ.var.to.zero=sum(circ.var.name%in%use)==0
		}else{
			set.circ.var.to.zero=F
		}
		usel=model.terms%in%use
		#if(length(use)>0)
		for(i in 1:length(model.terms)){
			if(is.factor(data[, model.terms[i]])){
				new.data[[i]]=levels(data[, model.terms[i]])
			}else if(!is.factor(data[, model.terms[i]]) & usel[i] & ifelse(length(circ.var.name)==0, T, !grepl(x=model.terms[i], pattern=circ.var.name))){
				new.data[[i]]=seq(from=min(data[, model.terms[i]]), to=max(data[, model.terms[i]]), length.out=resol)
			}else  if(!is.factor(data[, model.terms[i]]) & ifelse(length(circ.var.name)==0, T, !grepl(x=model.terms[i], pattern=circ.var.name))){
				new.data[[i]]=mean(data[, model.terms[i]])
			}
		}
		names(new.data)=model.terms
		if(length(circ.var.name)==1){
			new.data=new.data[!(model.terms%in%paste(c("sin(", "cos("), circ.var.name, ")", sep=""))]
			if(sum(grepl(pattern=circ.var.name, x=use))>0){
				new.data=c(new.data, list(seq(min(circ.var, na.rm=T), max(circ.var, na.rm=T), length.out=resol)))
				names(new.data)[length(new.data)]=circ.var.name
			}else{
				new.data=c(new.data, list(0))
			}
			model.terms=model.terms[!(model.terms%in%paste(c("sin(", "cos("), circ.var.name, ")", sep=""))]
		}
		xnames=names(new.data)
		#browser()
		new.data=data.frame(expand.grid(new.data))
		names(new.data)=xnames
		#browser()
		if(length(circ.var.name)==1){
			names(new.data)[ncol(new.data)]=circ.var.name
		}
		if(set.all.effects.2.zero){
			for(iterm in setdiff(colnames(new.data), c("(Intercept)", use))){
				new.data[, iterm]=0
			}
		}
		##prepare model frame for conditional model for prediction:
		model=attr(terms(as.formula(xcall[["cond.form"]])), "term.labels")
		model=model[!grepl(x=model, pattern="|", fixed=T)]
		cond.m.mat=model.matrix(object=as.formula(paste(c("~", paste(model, collapse="+")), collapse="")), data=new.data)
		if(set.circ.var.to.zero){
			cond.m.mat[,paste(c("sin(", circ.var.name, ")"), collapse="")]=0
			cond.m.mat[,paste(c("cos(", circ.var.name, ")"), collapse="")]=0
		}
		##get bootstrapped fitted values for the conditional part:
		ests=all.ind.boots$fe[, substr(colnames(all.ind.boots$fe), start=1, stop=5)=="cond@", drop=F]
		est.names=gsub(colnames(ests), pattern="cond@", replacement="")
		ci.cond=lapply(1:nrow(ests), function(x){
			return(cond.m.mat[, est.names, drop=F]%*%as.vector(ests[x, ]))
		})
		ci.cond=matrix(unlist(ci.cond), ncol=nboots, byrow=F)
		ci.cond=t(apply(ci.cond, 1, quantile, prob=c((1-level)/2, 1-(1-level)/2), na.rm=T))
		colnames(ci.cond)=c("lower.cl", "upper.cl")
		##get fitted value for the conditional part:
		ests=ci.estimates$fe
		ests=ests[substr(rownames(ests), start=1, stop=5)=="cond@", , drop=F]
		est.names=gsub(rownames(ests), pattern="cond@", replacement="")
		fv.cond=cond.m.mat[, est.names, drop=F]%*%ests$orig
		if(summary(model.res)$link=="log"){
			ci.fitted.cond=data.frame(fitted=exp(fv.cond), exp(ci.cond))
		}else if(summary(model.res)$link=="logit"){
			ci.fitted.cond=data.frame(fitted=plogis(fv.cond), plogis(ci.cond))
		}else if(summary(model.res)$link=="identity"){
			ci.fitted.cond=data.frame(fitted=fv.cond, ci.cond)
		}
		ci.fitted.cond=data.frame(new.data, ci.fitted.cond)
		
		##prepare model frame for ZI model for prediction:
		if(xcall[["zi.form"]]!="~0"){
			model=attr(terms(as.formula(xcall[["zi.form"]])), "term.labels")
			model=model[!grepl(x=model, pattern="|", fixed=T)]
			zi.m.mat=model.matrix(object=as.formula(paste(c("~", paste(model, collapse="+")), collapse="")), data=new.data)
			##get bootstrapped fitted values for the conditional part:
			ests=all.ind.boots$fe[, substr(colnames(all.ind.boots$fe), start=1, stop=3)=="zi@", drop=F]
			est.names=gsub(colnames(ests), pattern="zi@", replacement="")
			ci.zi=lapply(1:nrow(ests), function(x){
				return(zi.m.mat[, est.names, drop=F]%*%as.vector(ests[x, ]))
			})
			ci.zi=matrix(unlist(ci.zi), ncol=nboots, byrow=F)
			ci.zi=t(apply(ci.zi, 1, quantile, prob=c((1-level)/2, 1-(1-level)/2), na.rm=T))
			colnames(ci.zi)=c("lower.cl", "upper.cl")
			##get fitted value for the conditional part:
			ests=ci.estimates$fe
			ests=ests[substr(rownames(ests), start=1, stop=3)=="zi@", , drop=F]
			est.names=gsub(rownames(ests), pattern="zi@", replacement="")
			fv.zi=zi.m.mat[, est.names, drop=F]%*%ests$orig
			ci.fitted.zi=data.frame(fitted=plogis(fv.zi), plogis(ci.zi))
			#ci.fitted.zi=ci.fitted*(1-ci.fitted.zi)
			ci.fitted.zi=data.frame(new.data, ci.fitted.zi)
		}

		##prepare model frame for disp model for prediction:
		if(xcall[["disp.form"]]!="~0"){
			model=attr(terms(as.formula(xcall[["disp.form"]])), "term.labels")
			model=model[!grepl(x=model, pattern="|", fixed=T)]
			disp.m.mat=model.matrix(object=as.formula(paste(c("~", paste(model, collapse="+")), collapse="")), data=new.data)
			##get bootstrapped fitted values for the conditional part:
			ests=all.ind.boots$fe[, substr(colnames(all.ind.boots$fe), start=1, stop=5)=="disp@", drop=F]
			est.names=gsub(colnames(ests), pattern="disp@", replacement="")
			ci.disp=lapply(1:nrow(ests), function(x){
				return(disp.m.mat[, est.names, drop=F]%*%as.vector(ests[x, ]))
			})
			ci.disp=matrix(unlist(ci.disp), ncol=nboots, byrow=F)
			ci.disp=t(apply(ci.disp, 1, quantile, prob=c((1-level)/2, 1-(1-level)/2), na.rm=T))
			colnames(ci.disp)=c("lower.cl", "upper.cl")
			##get fitted value for the dispersion part:
			ests=ci.estimates$fe
			ests=ests[substr(rownames(ests), start=1, stop=5)=="disp@", , drop=F]
			est.names=gsub(rownames(ests), pattern="disp@", replacement="")
			fv.disp=disp.m.mat[, est.names, drop=F]%*%ests$orig
			ci.fitted.disp=data.frame(fitted=exp(fv.disp), exp(ci.disp))
			#ci.fitted.disp=ci.fitted*(1-ci.fitted.disp)
			ci.fitted.disp=data.frame(new.data, ci.fitted.disp)
		}
		ci.fitted=list(
			ci.fitted.cond=ci.fitted.cond, 
			ci.fitted.zi=ci.fitted.zi,
			ci.fitted.disp=ci.fitted.disp
		)
			
	}else{
		ci.fitted=NULL
	}
	return(list(ci.estimates=ci.estimates, ci.fitted=ci.fitted, all.boots=all.ind.boots))
}
```
