## Supplementary material for "Asian elephants (*Elephas maximus*) recognise human visual attention from body and face orientation": Data repository: glmmTMB_stability.r

```
glmmTMB.stab<-function(model.res, contr=NULL, ind.cases=F, para=F, data=NULL, use=NULL, n.cores=c("all-1", "all"), save.path=NULL, load.lib=T, lib.loc=.libPaths()){
	print("please carefully evaluate whether the result makes sense, and if not, please contact me")
	#function determining stability of GLMMs (run using lmer or glmer) by excluding levels of random effects, one at a time;
	#supports
		#weights, offset terms and random slopes;
	#does not support
		#correlations between random intercepts and random slopes
		#any terms calculated in the model formula (e.g., log, function I, etc.); but interactions do work
	#latest additions/modifications:
		#copes also with data sets where a level of a (fixed effects) factor is entirely dropped from the data
		#new way of catching warnings (are contained in the detailed output table)
		#includes sample size in the detailed output table
		#catches errors
		#use: an argument taking the names of the random effects for which model stability should be evaluated
			#(useful for models with a random effect having a unique case for each level of the data)
	#written by Roger Mundry
	#modified April 2016 (added dealing with glmer.nb; fixed a bug in the output of the random effects of tthe original model)
	#last modified Mar 2017 (added dealing with fixed effects factor levels dropped when dropping levels of random effects)
	#last modified June 14 2017 (fixed dealing with random effects comprising correlation parameters)
	#last modified July 06 2017 (fixed small bug happening when model revealed an error)
	#last modified Nov 26 2019 (copes with models uncluding no randm effects)
	if(load.lib){library(glmmTMB, lib.loc=lib.loc)}
	n.cores=n.cores[1]
	xx=as.character(model.res$call)
	names(xx)=names(model.res$call)
  model.fe.re=as.formula(xx["formula"])
  model.zi=as.formula(xx["ziformula"])
	model.disp=as.formula(xx["dispformula"])

	xcall=as.character(model.res$call)
	names(xcall)=names(model.res$call)
	xcall=list(cond.form=xcall["formula"], zi.form=xcall["ziformula"], disp.form=xcall["dispformula"], xfam=xcall["family"])
	if(grepl(xcall[["xfam"]], pattern="(", fixed=T)){
		xfam=xcall[["xfam"]]
		xfam=unlist(strsplit(xfam, split="(", fixed=T))
		xfam[2]=gsub(x=xfam[2], pattern=")", replacement="", fixed=T)
		xfam[2]=gsub(x=xfam[2], pattern="link = ", replacement="", fixed=T)
		xfam[2]=gsub(x=xfam[2], pattern="\"", replacement="", fixed=T)
		if(substr(xcall[["xfam"]], start=1, stop=5)!="Gamma"){
			xcall[["xfam"]]=get(xfam[1])(xfam[2])
		}else{
			if(xfam[2]=="log"){
				xcall[["xfam"]]=Gamma(link="log")
			}else if(xfam[2]=="inverse"){
				xcall[["xfam"]]=Gamma(link="inverse")
			}else if(xfam[2]=="identity"){
				xcall[["xfam"]]=Gamma(link="identity")
			}else{
				stop("Error: family not supported")
			}
		}
	}
	xfam=xcall[["xfam"]]
# 	
# 	xfam=xx["family"]
# 	if(grepl(xfam, pattern="(", fixed=T)){
# 		#xfam=xcall[["xfam"]]
# 		xfam=unlist(strsplit(xfam, split="(", fixed=T))
# 		xfam[2]=gsub(x=xfam[2], pattern=")", replacement="", fixed=T)
# 		xfam[2]=gsub(x=xfam[2], pattern="link = ", replacement="", fixed=T)
# 		xfam[2]=gsub(x=xfam[2], pattern="\"", replacement="", fixed=T)
# 		if(substr(xfam, start=1, stop=5)!="Gamma"){
# 			xfam=get(xfam[1])(xfam[2])
# 		}else{
# 			if(xfam[2]=="log"){
# 				xfam=Gamma(link="log")
# 			}else if(xfam[2]=="inverse"){
# 				xfam=Gamma(link="inverse")
# 			}else if(xfam[2]=="identity"){
# 				xfam=Gamma(link="identity")
# 			}else{
# 				stop("Error: family not supported")
# 			}
# 		}
# 		#xfam=get(xfam[1])(xfam[2])
# 	}
	##need to address how weights are recognized
	if(any(names(xx)=="weights")){
		data$XXXweights=data[, xx["weights"]]
	}else{
		data$XXXweights=1
	}
  ranefs=unique(unlist(lapply(ranef(model.res), names)))
	if(length(use)==0){use=ranefs}
	ranefs=intersect(ranefs, use)
  xlevels=lapply(ranefs, function(x){return(as.vector(unique(data[ ,x])))})
  ranefs=rep(ranefs, unlist(lapply(xlevels, length)))
  to.do=cbind(ranefs, unlist(xlevels))
  if(ind.cases){
    data=data.frame(data, ic=as.factor(1:nrow(data)))
    to.do=rbind(to.do, cbind("ic", levels(data$ic)))
  }
	extract.ranef.from.glmmTB<-function(m){
		#to.do=VarCorr(model.res)
		#varcor.names=names(to.do)
		to.do=summary(m)$varcor
		xx=lapply(to.do, function(x){
			lapply(x, attr, "stddev")
		})
		res=data.frame(
			what=rep(names(to.do), unlist(lapply(xx, function(x){sum(unlist(lapply(x, length)))}))),
			Groups=rep(unlist(lapply(to.do, names)), unlist(lapply(xx, function(x){unlist(lapply(x, length))}))),
			var1=unlist(lapply(xx, function(x){unlist(lapply(x, names))})),
			var2=NA,
			sdcor=unlist(xx)
		)
		xx=lapply(to.do, function(x){
			lapply(x, attr, "correlation")
		})
		xx=xx[unlist(lapply(xx, length))>0]
		xx=data.frame(
			what=rep(names(xx), unlist(sum(unlist(lapply(xx, function(x){lapply(x, function(x){prod(dim(x))})}))))),
			Groups=rep(unlist(lapply(to.do, names)), unlist(lapply(xx, function(x){lapply(x, function(x){prod(dim(x))})}))),
			var1=unlist(lapply(xx, function(x){lapply(x, function(x){rep(rownames(x), times=ncol(x))})})),
			var2=unlist(lapply(xx, function(x){lapply(x, function(x){rep(colnames(x), each=nrow(x))})})),
			sdcor=unlist(xx)
		)
		xx=xx[as.character(xx[, "var1"])<as.character(xx[, "var2"]), ]
		res=rbind(res, xx)
		res=res[order(as.character(res$what), as.character(res$Groups), as.character(res$var1)), ]
		return(res)
	}
	if(length(contr)==0){
		contr=glmmTMBControl()
	}
	n.ranef=nrow(extract.ranef.from.glmmTB(model.res))
  ifun<-function(x, model.res, to.do, data, contr, extract.ranef.from.glmmTB, n.ranef, xfam){
    sel.ii.data=data[data[,to.do[x, 1]]!=to.do[x, 2], ]
		sel.ii.res=try(glmmTMB(formula=model.fe.re, ziformula=model.zi, dispformula=model.disp, family=xfam, data=sel.ii.data, weights=XXXweights, control=contr), silent=T)
		#browser()
		if(length(save.path)>0){
			if(class(sel.ii.res)!="try-error"){
				est.fixed.effects=fixef(sel.ii.res)
				est.random.effects=summary(sel.ii.res)$varcor
				n=sel.ii.res$modelInfo$nobs
				what=to.do[x, ]
				converged=sel.ii.res$sdr$pdHess
				save(file=paste(c(paste(c(paste(c(save.path, "m"), collapse="/"), x), collapse="_"), ".RData"), collapse=""), 
					list=c("what", "est.fixed.effects", "est.random.effects", "n", "converged"))
			}else{
				ires="model revealed an error"
				save(file=paste(c(paste(c(paste(c(save.path, "m"), collapse="/"), x), collapse="_"), ".RData"), collapse=""), list=c("ires"))
			}
		}
    if(class(sel.ii.res)!="try-error"){
			fe=fixef(sel.ii.res)
			xnames=paste(rep(names(fe), times=unlist(lapply(fe, length))), unlist(lapply(fe, names)), sep="@")
			fe=unlist(fe)
			names(fe)=xnames
			RE=NULL
			if(any(!unlist(lapply(summary(sel.ii.res)$varcor, is.null)))){
				RE=extract.ranef.from.glmmTB(sel.ii.res)
				xx=apply(as.matrix(RE[, c("what", "Groups", "var1", "var2")]), 1, paste, collapse="@")
				RE=RE[, "sdcor"]
				names(RE)=xx
			}
			return(list(fere=c(fe, RE), N=sel.ii.res$modelInfo$nobs, converged=sel.ii.res$sdr$pdHess))
		}else{
      return(list(fere=rep(NA, sum(unlist(lapply(fixef(model.res), length)))+n.ranef), N=NA, converged="error"))
    }
  }
  if(para){
		#on.exit(expr = parLapply(cl=cl, X=1:length(cl), fun=function(x){rm(list=ls())}), add = FALSE)
		#on.exit(expr = stopCluster(cl), add = T)
    require(parallel)
    cl <- makeCluster(getOption("cl.cores", detectCores()))
		if(n.cores!="all"){
			if(n.cores=="all-1"){n.cores=length(cl)-1}
			if(n.cores<length(cl)){
				cl=cl[1:n.cores]
			}
		}
    parLapply(cl=cl, 1:length(cl), fun=function(x){
      library(glmmTMB, lib.loc=lib.loc)
      return(invisible(""))
    })
    all.coeffs=parLapply(cl=cl, X=1:nrow(to.do), fun=ifun, model.res=model.res, to.do=to.do, data=data, contr=contr, 
			extract.ranef.from.glmmTB=extract.ranef.from.glmmTB, n.ranef=n.ranef, xfam=xfam)
    parLapply(cl=cl, 1:length(cl), fun=function(x){
      return(rm(list=ls()))
    })
    stopCluster(cl=cl)
  }else{
    all.coeffs=lapply(1:nrow(to.do), ifun, model.res=model.res, to.do=to.do, data=data, contr=contr, 
			extract.ranef.from.glmmTB=extract.ranef.from.glmmTB, n.ranef=n.ranef, xfam=xfam)
  }
	all.n=unlist(lapply(all.coeffs, function(x){x$N}))
	all.conv=unlist(lapply(all.coeffs, function(x){x$converged}))
	xnames=unique(unlist(lapply(all.coeffs, function(x){names(x$fere)})))
	all.coeff.mat=matrix(NA, ncol=length(xnames), nrow=length(all.coeffs))
	colnames(all.coeff.mat)=xnames
	for(i in 1:length(all.coeffs)){
		all.coeff.mat[i, names(all.coeffs[[i]]$fere)]=as.vector(all.coeffs[[i]]$fere)
	}
	#extract results for original model:
	orig=fixef(model.res)
	xnames=paste(rep(names(orig), times=unlist(lapply(orig, length))), unlist(lapply(orig, names)), sep="@")
	orig=unlist(orig)
	names(orig)=xnames
	if(any(!unlist(lapply(summary(model.res)$varcor, is.null)))){	
		RE=extract.ranef.from.glmmTB(model.res)
		xx=apply(as.matrix(RE[, c("what", "Groups", "var1", "var2")]), 1, paste, collapse="@")
		RE=RE[, "sdcor"]
		names(RE)=xx
		orig=c(orig, RE)
	}
	#browser()
  xsum=apply(all.coeff.mat, 2, range, na.rm=T)
  xsum=data.frame(what=colnames(all.coeff.mat), orig=orig[colnames(all.coeff.mat)], t(xsum))
	rownames(xsum)=as.character(xsum$what)
  colnames(to.do)=c("ranef", "level")
  xx=apply(is.na(all.coeff.mat), 1, sum)
  if(sum(xx>0 & xx<ncol(all.coeff.mat))>0){
		warning(paste(c("for", sum(xx>0 & xx<ncol(all.coeff.mat)), "subset(s) the full model could not be fitted because of fixed effects factor levels dropped from the data"), collapse=" "))
	}
  all.coeff.mat=data.frame(to.do, N=all.n, converged=all.conv, all.coeff.mat)
  names(xsum)[3:4]=c("min", "max")
  return(list(detailed=all.coeff.mat, summary=xsum))
}
```
