## Supplementary material for "Asian elephants (*Elephas maximus*) recognise human visual attention from body and face orientation": Data repository: HLJ_readme.txt

This repository contains the data and R code for the manuscript:
"Asian elephants (Elephas maximus) recognise human visual attention from body and face orientation"

Authors: Hoi-Lam Jim, Shinya Yamamoto, Pakkanut Bansiddhi, Joshua M. Plotnik

Description of the data and file structure

This study investigates whether Asian elephants (Elephas maximus) are sensitive to the visual perspective of a human. We conducted a similar study to Smet and Byrne (2014) to test whether Asian elephants modify the frequency of experimenter-directed signals in a food-requesting task according to whether the experimenter's body and/or face is oriented towards them.

Folder structure and contents

analysis/

Contains the R Markdown file used for the statistical analyses: ele_vis_att_analysis.rmd

data/

Contains the datasets used for the analyses and inter-rater reliability:

ele_vis_att_data.csv: Main dataset used in the analysis.

Variables:

subject (character): Elephant's name (unique identifier for each subject).

session (numeric): Session number (1–4), repeated measures across sessions for each subject.

trial (numeric): Trial number (1–5), repeated measures within each session.

condition (factor with 5 levels, within-subjects): Experimental condition. The levels are:

Ba_Fa: Body away, face away

Ba_Ft: Body away, face towards

Bt_Fa: Body towards, face away

Bt_Ft: Body towards, face towards

Np: Not present

Behavioural variables: All behavioural variables are count variables. Detailed descriptions are provided in Table 1 of the manuscript. These include: 

trunkswing

trunkout_forward

trunkout_self

trunkout_ground

trunkup_forward

trunkup_self

trunkup_other

reachout_experimenter

reachout_food

trainedbegging

headnod

totalbehaviours (integer): Total number of behaviours coded during the trial.

SV_coding.csv: Raw behavioural coding by the research assistant (SV) for inter-rater reliability checks. Follows the same format as ele_vis_att_data.csv

interreliability_data.csv: Dataset with two columns (HLJ and SV's codings) used to compute the Intraclass Correlation Coefficient (ICC). The totalbehaviours column was extracted from HLJ’s and SV’s datasets.

functions/

Contains R functions provided by Mundry (2023) for model stability checks, confidence interval calculations, and overdispersion tests.

graphics/

Contains the plot used in the manuscript:

fig3: Figure 3 in the manuscript.

Code/software

All statistical analyses were conducted using R (v4.4.1) in RStudio (v2024.04.2+764). All R packages used are referenced in the manuscript and listed in the R Markdown file (ele_vis_att_analysis.rmd) located in the analysis/ folder. All R functions provided by Mundry (2023) used are included in the functions/ folder. 

Additional information

This study was pre-registered: https://aspredicted.org/J32_HR5

References (in this ReadMe)

Mundry R. 2023 some R functions. Zenodo. https://doi.org/10.5281/zenodo.7670524

Smet AF, Byrne RW. 2014 African elephants (Loxodonta africana) recognize visual attention from face and body orientation. Biol. Lett. 10, 20140428.
