## Supplementary figures and images for "Asian elephants (*Elephas maximus*) recognise human visual attention from body and face orientation"

### fig3.png

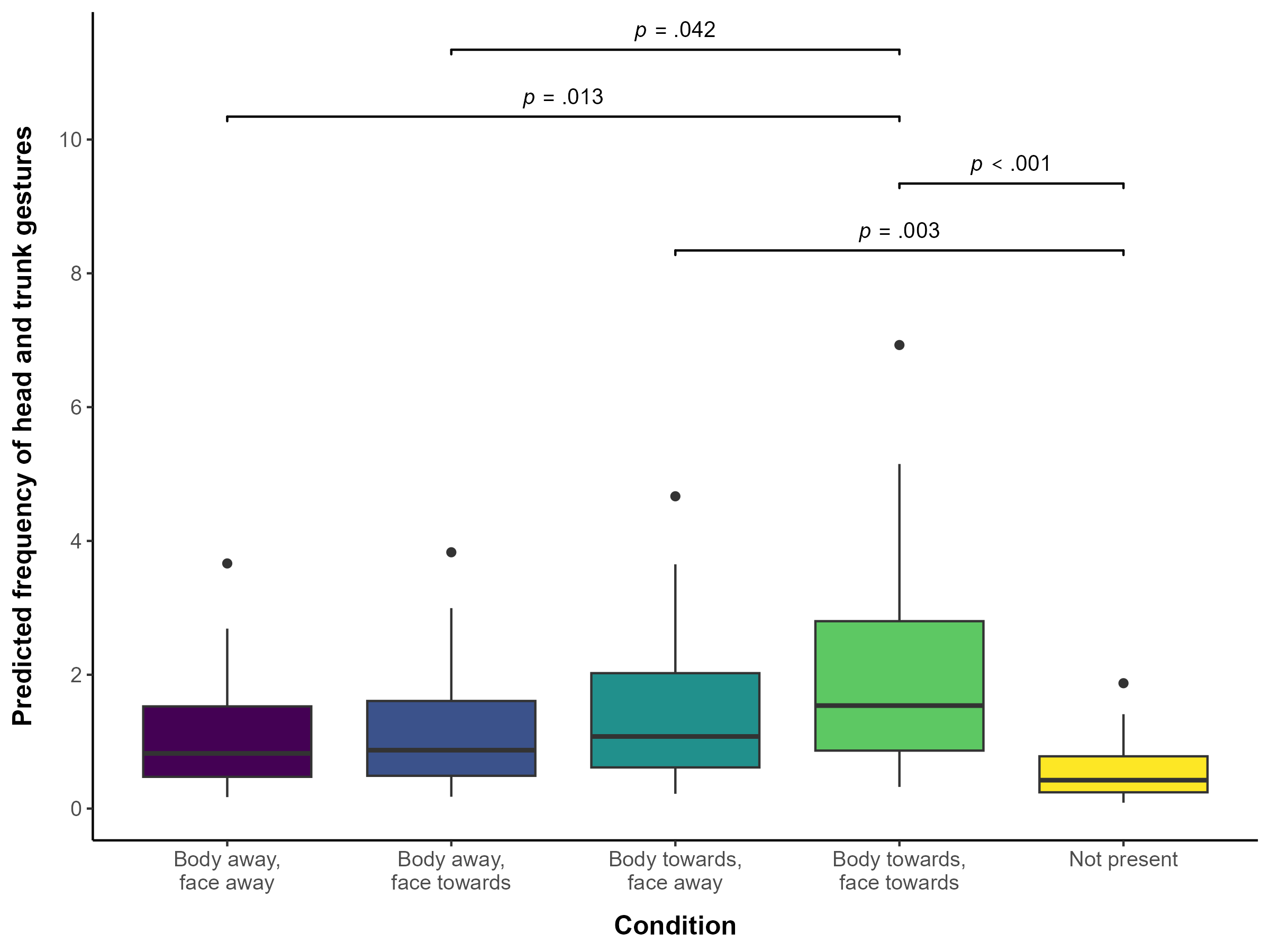
